# Alpha-synuclein fibrils induce budding of mitochondrial-derived vesicles

**DOI:** 10.1101/2025.10.13.682220

**Authors:** Thomas Braun, Cinzia Tiberi, Viviane Reber, Dhiman Ghosh, Roland Riek, Tetiana Serdiuk

**Author notes:** Corresponding authors: Tetiana Serdiuk, Thomas Braun. equal contribution (Cinzia Tiberi, Viviane Reber). **Competing Interest Statement:** The authors declare no competing interests.

## Abstract

α-synuclein (α-syn) aggregation is a hallmark of synucleinopathies, a class of neurodegenerative disorders such as Parkinson’s disease (PD). Several lines of evidence indicate the involvement of mitochondria in the disease pathology. Despite extensive study, the link between α-syn aggregation and mechanisms of mitochondrial toxicity remains not fully understood. Using high-resolution imaging with electron microscopy, we examined cells exposed to α-syn fibrils vs control cells with a focus on mitochondria. We found that upon exposure to α-syn fibrils, mitochondria increase in size, cristae structure gets defects, and mitochondria enhance the budding of mitochondrial-derived vesicles (MDVs). MDV formation reflects an evolutionarily conserved mechanism reminiscent of bacterial outer membrane vesicle (OMV) biogenesis. Structural proteomics analysis by mass spectrometry corroborates this microscopy observation by identifying changes in multiple proteins that regulate cristae structure, MDV formation, and trafficking. Our results provide a new link between α-syn and mitochondria and identify novel pathways responding to α-syn aggregates, particularly that α-synuclein directly triggers MDV generation. The processes we detected could be of interest for diagnostics and potential therapeutic interventions.

## Introduction

Both α-synuclein (α-syn) and mitochondria contribute to multiple neurodegenerative diseases collectively called synucleinopathies, particularly Parkinson’s (PD). The aggregation and pathological accumulation of α-syn are hallmarks of PD, where mitochondrial dysfunction is seen as a contributor and a consequence of the α-syn pathology^1,2^. Furthermore, environmental factors such as toxins like rotenone and MPTP induce PD-like pathology by impairing mitochondrial respiratory chain function, promoting oxidative stress, and triggering α-syn aggregation.^3-5^

α-syn aggregation into amyloid fibrils, which then deposit as inclusions called Lewy bodies, is PD’s hallmark. Recent ultrastructural analyses of Lewy bodies in PD patients’ brains have revealed that these pathological α-syn inclusions are often rimmed with damaged mitochondria.^6^ This suggests mitochondrial dysfunction in PD pathology. α-syn has shown a conformation-dependent influence on mitochondrial dynamics and function^7^, including fusion, fission, ATP synthesis^8,9^, and transport along microtubules^10^. Still, our knowledge of how α-syn fibrils impact mitochondria and of the crosstalk between α-syn and mitochondria in pathology remains limited.

Mitochondria are large organelles with peculiar structures, and many different defects can be directly seen using high-resolution imaging methods such as electron microscopy (EM). Here, we set out to characterize how mitochondrial morphology changes in response to a cell’s internalization of α-syn fibrils and how mitochondrial proteomes respond to fibrillar invasion. We discovered previously unknown responses of mitochondria to α-syn; in particular, we found that internalization of α-syn fibrils caused cristae defects and enhanced mitochondrial-derived vesicle (MDVs) budding, linking α-syn toxicity to a recently discovered mitochondrial quality control system-MDV pathway.

## Results

We studied how α-syn fibrils impact mitochondria morphology using the well-established endocytic uptake of α-syn fibrillar fragments as a model. In short, we used fibrillar aggregates of purified full-length, N-terminally acetylated human α-syn, which were fragmented to fibril lengths of approximately 50 nm. We exposed SH-SY5Y cells to the α-syn fibrillar fragments (250 nM monomer equivalent concentration), for 24 h, washed them with cell culture media two times, and fixed them with a solution of 2% paraformaldehyde and 2.5% glutaraldehyde afterward. Fixed cells were resin-embedded, cut into slices, and imaged by EM (Methods). At least 10 cells were analyzed per condition from different preparations.

We have analyzed EM data on several different axes regarding mitochondrial morphology: size, cristae shape and budding of MDVs. A representative overview of non-seeded (A) and α-syn seeded (B) cells is given in Figure 1. Interestingly, we observed a significantly larger area and perimeter of mitochondria when fibrils were added (*n*=289 and *n*=303 for unseeded and seeded cells, respectively), indicating potential mitochondrial swelling caused by α-syn (Figure 1C, D).

**Figure 1.**
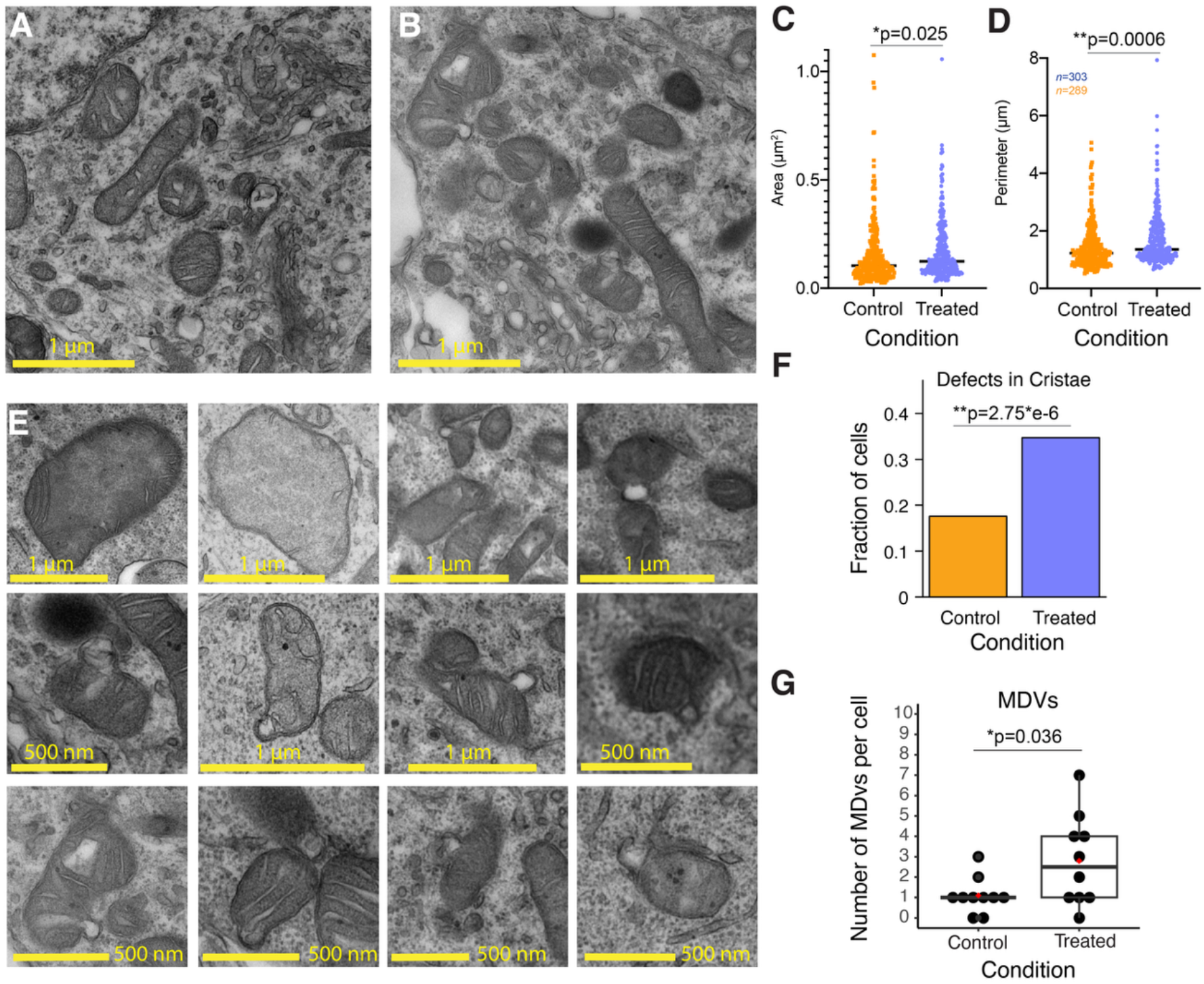
α-syn fibrils induce mitochondrial defects. **(A, B)** Representative EM images of mitochondria in control SH-SY5Y cells (**A**) and SH-SY5Y cells treated with α-syn fibrils (250 nM monomer equivalents) for 24h (**B**). (**C, D**) Distribution of the mitochondria area (**C**) and perimeter (**D**) in control cells and cells treated with α-syn fibrils. (**E**) Representative EM images showing cristae defects (first three images) and budding of mitochondrial derived vesicles. (**F**) Fraction of cells showing mitochondria with cristae defects. (**G**) Number of MDVs detected per cell for control and α-syn treated condition. P values were calculated using two tailed t test (**C, D, G**) and Fisher’s exact test (**F**). **p val< 0.01, *p val<0.05, NS p val >0.05. The medians are shown as black lines. The mean values (in **G**) are shown as red dots.

Representative ‘zoom-in’ images of mitochondria showing different morphological aberrations upon α-syn treatment are shown in Figure 1E. While inspecting cells upon treatment with α-syn fibrils, we found that mitochondria cristae significantly more often showed peculiar defects (Figure 1F): an atypical cristae shape (e.g., trapezia, triangle shape) or lack/depletion of cristae structure. Often, mitochondria depleted of cristae structure were also obviously larger than the other mitochondria. Close inspection of mitochondria in both conditions revealed differences in membrane budding. Several examples of mitochondria budding MDVs are shown in Figure 1E. MDVs are one of the pathways of mitochondrial quality control used by a cell to remove damaged material from mitochondria when mitophagy is still avoidable. We have seen approximately two times more MDVs on the EM images of cells treated with α-syn fibrils than the control cells treated with the same volume of PBS (Figure 1G). This result could indicate the response to enhanced mitochondrial stress caused by α-syn fibrils.

To support our visual results on the molecular level, we performed an unbiased structural proteomics analysis with Limited proteolysis coupled to mass spectrometry (LiP-MS). LiP-MS probes proteome accessibility to sequence unspecific protease, proteinase K (PK), across different conditions. LiP-MS can detect variable structural changes, including changes in protein-protein interactions, enzymatic activity, aggregation, conformational changes, and post-translational modifications^11^. We applied LiP-MS to cells treated with α-syn fibrils and control cells treated with PBS and found 130 proteins structurally altered upon treatment with α-syn (Figure 2A). Gene Ontology enrichment analysis for cellular components revealed enrichment of many mitochondria-related terms (Figure 2B), including mitochondria inner membrane and intermembrane space, MICOS complex, prohibitin complex, and MIB complexes. Multiple of these mitochondrial proteins are important for mitochondrial cristae organization. For instance, several proteins of the MICOS complex (Mic60, Mic19, Mic26) were among the strongest screening hits. The outer membrane translocase elements TOM40 and TOM70 and the inner membrane translocase TRIM23 changed the PK susceptibility upon α-syn treatment (Figure 2A). All these proteins are crucial for mitochondria cristae structure, mitochondrial biogenesis, maintenance of contact sites between inner and outer mitochondrial membrane, and mitochondria membrane remodeling. LiP-MS allowed us to identify the region of structural alteration on these proteins (Figure 2C). Remarkably, the Tollip protein, shown to be essential for TOM complex-positive MDV trafficking, along with the TOM complex, structurally responded to the α-syn fibrils treatment, corroborating our EM data on enhanced MDVs formation.^12^ Given its important function in maintaining mitochondria architecture, the MICOS complex was also suggested to play a role in the formation of MDVs^13^, possibly interacting with Rho GTPases MIRO1/MIRO2. Interestingly, along with detected structural changes in many sub-units of MICOS complex, we also detected a moderate structural change for MIRO2 (FC>1.5, p val<0.05).

**Figure 2.**
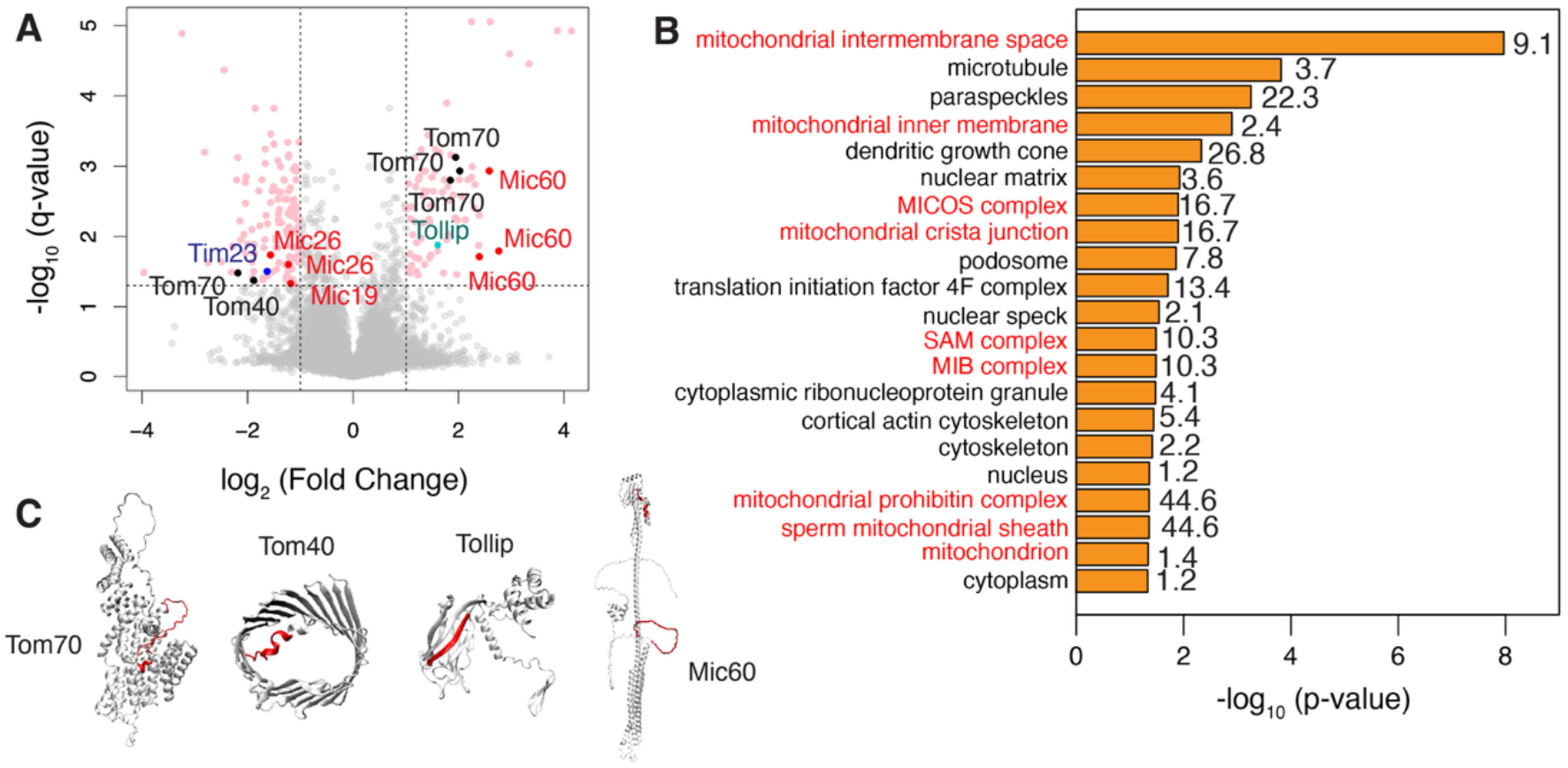
α-syn fibrils affect the mitochondrial structural proteome of living cells. (**A**) Volcano plot of proteins showing structural change due to uptake of α-syn fibrils. Significance cut-off of q val<0.05 and FC>2 was used. (**B**) Gene Ontology (Cellular component) enrichment analysis of LiP-MS hits. Fold enrichment is given by numbers for each enriched term. (**C**) Examples of LiP-MS hits with changing peptides mapped on the solved or predicted structures for TOM70 (PDB ID AF-O94826-F1), TOM40 (PDBID 7ck6), Tollip (AF-Q9H0E2), and MIC60 (AF-Q16891-F1).

## Discussion

Mitochondrial quality control is performed through various mechanisms, including the activity of mitochondrial proteases, mitophagy, and mitochondrial-derived vesicles (MDVs)^14^. The role of mitophagy in PD is better understood, even with genetic links to the disease being established (PINK1, Parkin). MDVs, a recently discovered quality control system in mitochondria, is proposed to be an alternative to mitophagy pathway^14^. MDVs selectively transport damaged mitochondrial components, including proteins and lipids, to lysosomes or peroxisomes for degradation. We found increased MDV generation upon cell treatment with α-syn fibril fragments. Furthermore, the mitochondria of the cells treated with α-syn fibrils were bigger and had more cristae defects than the control cells. Some large mitochondria were almost completely depleted of cristae.

In support of our microscopy data, structural proteomics revealed changes in multiple proteins important to the MDV pathway in response to α-syn treatment. For these and other LiP-MS mitochondrial protein hits, we mapped regions of structural alteration with a resolution of approximately 10 amino acids (Figure 2C).

The enhanced release of MDVs was recently described in murine brains with Down Syndrome^15^, and the findings were confirmed in human postmortem brain material. It has been suggested that MDV release is enhanced in Down Syndrome and Alzheimer’s disease and neurodegeneration in general^16^, and that MDVs are released extracellularly^15,16^. Therefore, MDVs are potentially interesting for early diagnostics. Here, we showed that α-syn fibrils can directly trigger enhancement of MDV release, establishing the link between α-syn aggregation and mitochondria stress. It remains to be evaluated whether tau and amyloid-β fibrils also can directly cause the enhancement of MDV formation.

## Methods

### Cell seeding

N-terminally acetylated α-syn was purified as described previously^17^. SH-SY5Y neuroblastoma cells (ATCC) were cultured in DMEM/F-12 GlutaMAX medium (#10565018, Thermo Fisher Scientific) supplemented with 10% fetal bovine serum (FBS) and 1% penicillin/streptomycin. For treatment, cells were exposed to 250 nM α-syn fibrils (concentration for monomer equivalent) in the complete culture medium for 24 hours. Mock-treated control cells were cultured in parallel under identical conditions.

### Electron microscopy

SH-SY5Y cells treated with αSyn fibrils or mock cells were fixed using a solution containing 2% paraformaldehyde (PFA) and 2.5% glutaraldehyde (GA) in 0.1M PIPES buffer. Following fixation, the cells were impregnated in 2% low-melting-point agarose and placed on ice for 20 minutes to allow the agarose to solidify. The resulting solid agarose pellets were carefully removed from the tubes and trimmed into smaller pieces using a razor blade.

Agarose cubes containing sample material were transferred to small glass vials, thoroughly washed with 0.1 M Cacodylate buffer, and post-fixed in 1% buffered osmium tetroxide for 1 hour at 4 °C. Samples were then rinsed with distilled water and stained *en bloc* with aqueous uranyl acetate for 1 hour at 4 °C in the dark. After staining, samples were dehydrated through a graded ethanol series. After three changes of absolute ethanol, samples are washed in acetone and finally embedded in a mixture of resin/acetone first and then in pure Epon 812 resin. Embedding is carried out in a 60°C oven for 48 h until the epoxy resin is completely hardened and ready to section.

Ultrathin sections (70 nm thick) were cut using an ultramicrotome (Leica EM UC7, Leica Microsystems, Austria) and contrasted with uranyl acetate and lead citrate. The sections were imaged using a FEI Tecnai G2 Spirit transmission electron microscope (TEM) operated at 80 kV. Images were acquired with anEMSIS Veleta digital camera (top, side-mounted) using RADIUS software (EMSIS GmbH). EM images were analyzed using Fiji. Two-tailed t test and Fisher’s exact test were used to access statistical significance.

### LiP-MS

### Proteomics analysis

#### Native cell lysis

Pellets of SH-SY5Y cells were resuspended in 200 μl of ice-cold LiP buffer (100 mM HEPES, 150 mM KCl, 1 mM MgCl_2_, pH 7.4). To lyse cells, we performed 10 cycles of 10 douncing steps using a pellet pestle on ice. A bicinchoninic acid (BCA) assay was used to measure protein concentrations. After native lysis and concentration determination, the samples were processed directly by LiP-MS.

#### LiP-MS

The concentrations of cell lysates were generally similar between replicates, with minor differences adjusted with LiP buffer (20 mM HEPES, 150 mM KCl, 1 mM MgCl_2_, at pH 7.4). All samples were diluted in LiP buffer to a final protein concentration of 2mg/ml and a total volume of 50 μl per sample. Six replicates per each condition were used.

LiP-MS experiments were conducted as described previously with some minor modifications.^11^ Briefly, Proteinase K (PK) was added at a 1:100 enzyme-to-substrate ratio in a total volume of 5 μl using a multi-channel pipette. Samples were briefly mixed by gentle pipetting up and down 5 times. Next, the samples were incubated at 37°C for exactly 5 min in a Biometra TRIO thermocycler. PK digestion of proteomes was stopped by incubating the samples at 99°C for 5 min, followed by a 5 min incubation on ice. Subsequently, 55 μl of freshly prepared 10% sodium deoxycholate (DOC) was added to each sample.

#### Trypsin / LysC digestion

Samples were reduced with tris(2-carboxyethyl)phosphine hydrochloride (TCEP) to a final concentration of 5 mM at 37 °C for 40 minutes with shaking at 800 rpm in a thermomixer (Eppendorf). Following reduction, samples were alkylated by adding iodoacetamide (IAA) to a final concentration of 40 mM and incubated for 30 minutes at room temperature in the dark. To reduce the concentration of sodium deoxycholate (DOC) to 1%, samples were diluted with 100 mM ammonium bicarbonate (Ambic). Proteolytic digestion was carried out by adding Lysyl endopeptidase (LysC, Wako Chemicals) and sequencing-grade trypsin (Promega) at an enzyme-to-substrate ratio of 1:100. Digestion was performed overnight at 37 °C in a thermomixer with continuous shaking at 800 rpm. The reaction was stopped by adding formic acid (FA, Carl Roth GmbH) to a final concentration of 3%, resulting in DOC precipitation. DOC was removed by three rounds of centrifugation at 21,000 rcf for 20 minutes each, with the supernatant collected after each spin. The combined supernatants were desalted using Sep-Pak tC18 cartridges (Waters) and eluted with 80% acetonitrile containing 0.1% formic acid.

### LC-MS/MS data acquisition

#### Liquid chromatography

LiP-MS samples were analyzed using an Orbitrap Exploris mass spectrometer (Thermo Fisher Scientific) coupled to a nano-electrospray ion source and a nano-flow LC system (Easy-nLC 1200, Thermo Fisher Scientific). Peptide separation was performed using self-packed 40 cm x 0.75 mm columns (New Objective) containing 1.9 μm C18 beads. A linear gradient of LC buffer A (5 % ACN, 0.1 % FA, Carl Roth GmbH) and LC buffer B (95 % ACN, 0.1 % FA, Carl Roth GmbH) increasing from 3 % to 30 % of LC buffer B over 120 minutes at a flow rate of 300 nl/minute was used.

#### Data-independent acquisition

Data-independent acquisition (DIA) scans were performed using 41 variable-width isolation windows. Precursor ions were isolated with a quadrupole. For MS1 survey scans, a mass range of 350-1150 m/z was used and an Orbitrap resolution of 120’000. A normalized automated gain control (AGC) target of 200 % was applied. High-energy collision induced dissociation (HCD) was employed to fragment precursor ions. DIA-MS/MS spectra were recorded in the Orbitrap with a resolution of 30’000 and a mass range of 350-1150 m/z. The maximal injection time was set to 66 ms.

#### Data-dependent acquisition

Samples were acquired in Data-dependent mode (DDA) alongside DIA runs to generate spectral libraries. Precursor ions were isolated using a quadrupole. For MS1 survey scans, a mass range of 350-1150 m/z was used. Precursor ions with intensities above 5’000 were selected for MS/MS scans. The isolation window of the quadrupole was 1.4 m/z. HCD was employed to fragment precursor ions. MS/MS spectra were acquired on an Orbitrap at a resolution of 30’000, an AGC target of 200% and a maximum injection time of 54 ms.

### Data search, statistical data analysis and visualisation

Spectral libraries were generated through a Pulsar search in Spectronaut (Biognosys AG). The raw DIA MS data were also searched in Spectronaut using generated libraries. Compared to the default settings, the specificity was adjusted to semi-specific for Trypsin/P. This adjustment accounts for semi-tryptic peptides generated by PK. LiP-MS data were analysed at the peptide level. To compare peptide abundance, we performed a two-tailed Welch t-test with Benjamini-Hochberg correction after median normalization. p-Values adjusted for multiple comparisons were referred to as q-values. Protein group quantities were used for protein abundance comparison. The significance threshold FC > 2, q-val < 0.05 was applied. The statistical analysis was conducted in R using the protti and limma packages. For normalization, peptide intensities were divided by the corresponding control protein abundance per peptide before comparing peptide abundances. Protein abundance was assessed in the same lysates that had been digested with trypsin only in parallel to the LiP-MS workflow. The same sample processing protocol was followed for tryptic control samples, except adding PK. GO enrichment analysis was carried out using David. The identified protein set served as the background for GO Enrichment analysis. LiP-MS hits were visualized and mapped on protein structures using Visual Molecular Dynamics (VMD).

## Acknowledgments

We thank Gelu foundation and Synapsis Foundation for funding this project (funded to T. S, CDA-2023-CDA02). We acknowledge support by the Swiss Nanoscience Institute (ARGOVIA FuncEM, SNI PhD 1901) and the Swiss National Science Foundation (projects 200021_162521, 200020_192190) granted to T.B. We thank Prof. Paola Picotti (ETHZ) for providing access to the mass spectrometry instruments.

